# Bulk and Single-cell Transcriptomic Brain Data Identify Overlapping Processes and Cell-types with Human AUD and Mammalian Models of Alcohol Use

**DOI:** 10.1101/2024.07.02.601528

**Authors:** Spencer B. Huggett, Sharmila Selveraj, John E. McGeary, Ami Ikeda, Emerald Yuan, Lauren B. Loeffel, Rohan H.C. Palmer

## Abstract

This study explores the neurobiological underpinnings of alcohol use disorder (AUD) by integrating bulk and single-cell transcriptomic data from humans, primates, and mice across three brain regions associated with addiction (i.e., prefrontal cortex (PFC), nucleus accumbens (NAc), and central amygdala (CeA)). We compared AUD RNA expression and cell-type abundance from 92 human brain to data from 53 primates and 90 mice engaged in diverse alcohol use paradigms. The findings revealed significant and reproducible correlations between human AUD and mammalian models of alcohol use that vary by tissue, species, and behavioral paradigm. The strongest correlations occurred between primate and mouse models of binge drinking (i.e., high drinking in the dark). Certain primate models demonstrated that the brain RNA correlations with human alcohol use disorder (AUD) were approximately 40% as strong as the correlations observed within human samples themselves. By integrating single-cell transcriptomic data, this study observed decreased oligodendrocyte proportions in the PFC and NAc of human AUD with similar trends in animal models. Gene co-expression network analyses revealed conserved systems associated with human AUD and animal models of heavy/binge alcohol consumption. Gene co-expression networks were enriched for pathways related to inflammation, myelination, and synaptic plasticity and the genes within them accounted for ∼20% of the heritability in human alcohol consumption. Identified hub genes were associated with relevant traits (e.g., impulsivity, motivation) in humans and mice. This study sheds light on conserved biological entities underlying AUD and chronic alcohol use, providing insights into the cellular, genetic, and neuromolecular basis across species.

## Introduction

Substantial progress has been made in our biological understanding of alcohol use disorder (AUD). AUD is influenced by a constellation of genetic factors ^1^ as well as multiple tissues and cell types predominantly in the brain ^2^. Scant evidence exists on whether, or how the molecular and cellular findings from animal models of heavy alcohol use map onto the neurobiological processes of human AUD ^3^. A deeper understanding of how human AUD data correlate with pre-clinical alcohol use paradigms could pinpoint conserved biological entities of AUD and chronic alcohol use.

A rich array of mammalian alcohol use models are utilized for investigating specific bio-behavioral components of human AUD ranging from chronic ethanol exposure ^4^ self-administration ^5^ and binge drinking ^6^. Various species and strains of animals are used that differ in the degree of volitional alcohol consumption and may differ in how they model human AUD.

Multi-omic datasets on human and animal alcohol use traits afford the opportunity to systematically explore the resemblance, or distinctions, of molecular patterns of alcohol use across species, tissues, and behavioral models. Most publicly available datasets include brain RNA data in the reward and anti-reward pathways in the brain that govern various addictive behaviors. The cellular make-up, neuroanatomical organization, and molecular features (i.e., RNA expression) in the brain are fairly conserved across species (^7^; ^8^; ^9^). To our knowledge, no study has investigated how AUD- and brain-region specific RNA expression and cell-types are conserved across non-human primate and mouse models of alcohol use.

Our study systematically compares pre-clinical paradigms of alcohol use and their relevance to human AUD. To identify conserved neurobiological entities underlying chronic alcohol use, we integrated bulk- and single-cell bioinformatics data from the three brain regions important for addiction in humans, primates, and mice. We hypothesized that human AUD RNA expression and cell type abundance would show consilience with animal models of alcohol use, but that the degree and nature of conservation would depend on 1) species, 2) brain region, and 3) behavioral paradigm. Specifically, we predicted that biological correlates of human AUD would overlap most with 1) primate models of alcohol use, 2) within limbic brain regions, and 3) in behavioral paradigms of binge drinking. Lastly, we predicted that genes associated with AUD and alcohol use across species would be enriched for genes that comprise the heritability of alcohol use traits.

## Methods

### Samples

Clinical and preclinical transcriptomic brain data on alcohol use was used in this paper. In total, we used bulk RNA-seq and microarray data on 92 humans, 53 primates, and 90 mice (see Table 1).

**Table 1.**
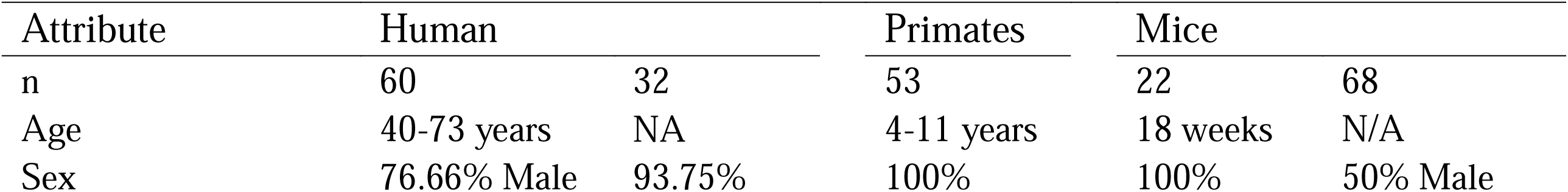
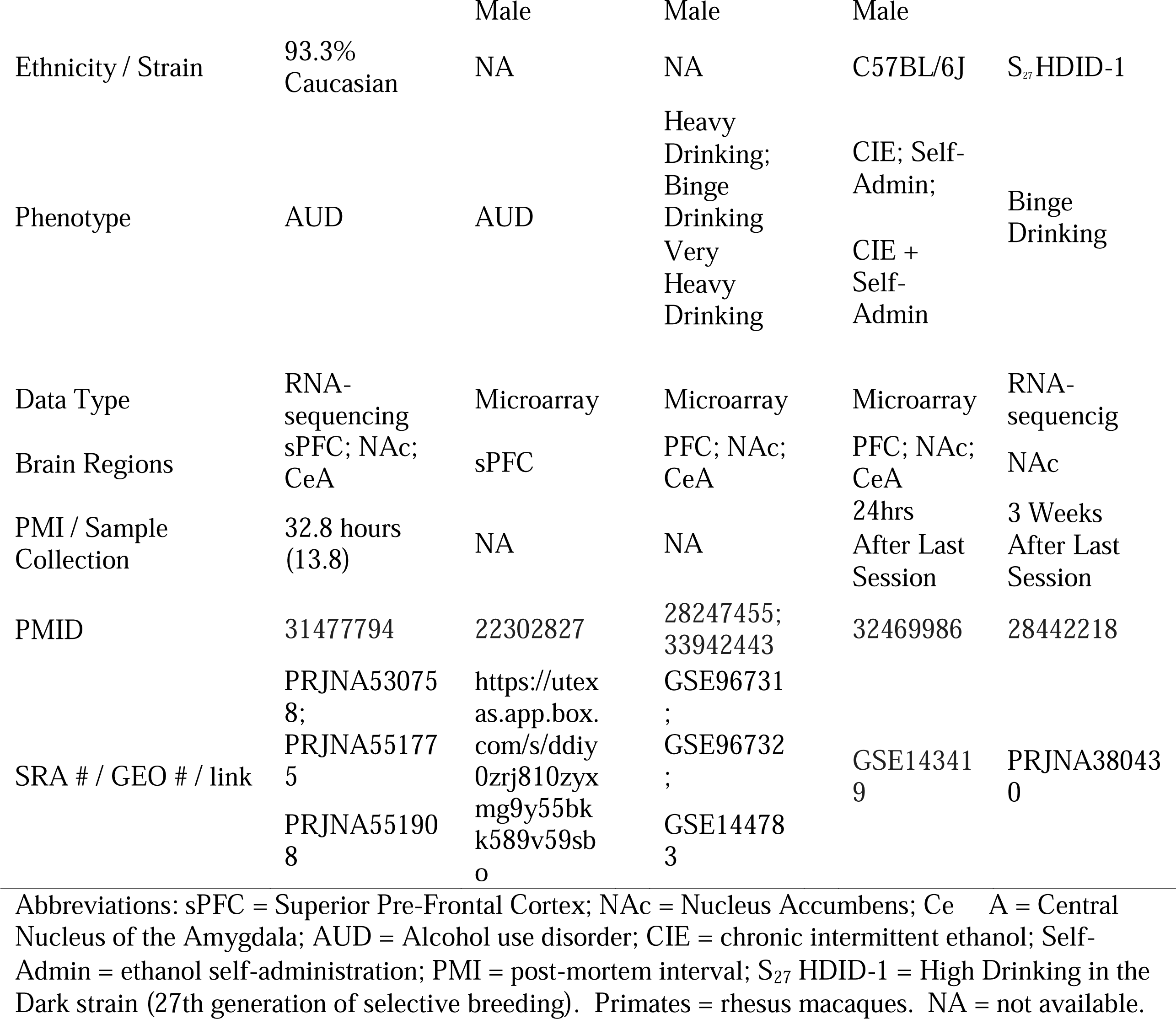
Descriptive Information on the Chronic Alcohol Use Samples Used in the Study.

### Human Transcriptomic Data

Two human AUD post-mortem brain samples were used – a discovery and replication sample. The discovery dataset included RNA-sequencing (RNA-seq) samples from the New South Wales Brain Tissue Resource Center. We acquired RNA-seq data from superior prefrontal cortex (PFC), nucleus accumbens (NAc), and central nucleus of the amygdala (CeA; ^10^). These samples were collected within three days of death from 30 individuals diagnosed with AUD and 30 social/non-drinking controls. The human replication sample used microarray data from the same resource using superior PFC tissue in an independent sample of 17 individuals with AUD and 15 matched controls ^11^.

### Primate Transcriptomic Data

Various preclinical alcohol use models were used that incorporated wide-ranging behavioral models, and a couple of mammalian species that used transcriptomic data in similar brain regions. Male primate (Rhesus macaque) RNA-seq data stemmed from four cohorts in the Monkey Alcohol Tissue Research Resource, originating from two separate studies. The first study used primate brain data from the PFC (cortical area 32) and the CeA ^12^. The second primate study focused on NAc core ^13^. Primates had four months of schedule-induced polydipsia to provoke daily drinking. After induction, primates underwent two 6-month sessions of alcohol self-administration where alcohol was freely available for over 90% of the day. Prior literature has demonstrated that primate alcohol consumption is comparable to humans with AUD within this paradigm ^14^. Primates show different levels of alcohol consumption and were split into multiple groups, including controls (alcohol naïve; NAc only), low drinkers, high drinkers, very high drinkers (ethanol consumption > 3g/kg for > 20% of days), or binge drinkers (CeA and PFC only). In the NAc, drinking groups were compared with alcohol naïve controls. For CeA and PFC data, drinking groups were compared with the low drinkers (PFC and CeA – as no alcohol naive controls existed in these data). After outlier removal (due to low RNA-seq read count; z-score < -2 of scale factor of normalized read counts), we used a total of 83 monkey brain samples (nNAc = 23; nCeA = 30; nPFC = 30).

### Mice Transcriptomic Data

Two studies investigated four drinking traits within mouse models. One used the high drinking in the dark (HDID) binge drinking paradigm in 68 male and female genetically diverse mice that were bred for drinking or controls (Heterogenous Stock-Northport mice; ^15^). The HDID binge drinking mouse sample used RNA-seq of NAc tissue. The drinking in the dark paradigm starts three hours into the dark phase and includes a four-day self-administration paradigm (20% ethanol) with the first three days consisting of three separate 2-hour sessions and the fourth day consisting of a single 4-hour session. Mice were sacrificed three weeks after their last session. The second mouse sample leveraged summary data from a study on adult male C57BL/6J mice (n = 8-11 per group; ^16^) assessing alcohol self-administration, chronic intermittent ethanol exposure (CIE), and the combination of CIE and alcohol self-administration. Alcohol self-administration involved a two-week acclimation followed by six weeks of a two-hour, two-bottle choice paradigm with 15% ethanol or non-alcoholic solution. CIE, modeling non-voluntary effects of chronic alcohol exposure, consisted of inhaling ethanol or air through a vapor chamber for 16 hours/day over four days, followed by a three-day abstinence period, totaling four cycles. CIE mice typically exhibit blood ethanol concentrations of 175-225 mg/dL ^17^. Combined CIE and alcohol self-administration involved a two-week acclimation, followed by alcohol self-administration for five days after each CIE cycle, capturing an escalated drinking phenotype. Mouse brains were harvested 24 hours after the last session, with ethanol-naïve mice serving as controls for differential expressions.

### Single-cell Transcriptomic Data

Single-nucleus and single-cell RNA sequencing data (sn/scRNA-seq), provided by 10x Genomics, were utilized from three studies involving humans ^18^, primates ^19^, and mice ^20^. These studies analyzed brain regions corresponding to those examined in the bulk transcriptomic samples. Human brain regions included the dorsal-lateral PFC (n=3), NAc (n=8), and amygdala (n=5). Mid-fetal primate scRNA-seq data included dorsal-lateral PFC (n = 2), striatum (n = 2), and amygdala (n = 2). Lastly, young adult male (8-10-week-old) C57BL/6J mice included the NAc (n = 11). The original cell-type identities were used from the original publications, but data underwent further quality control filtering out cells/nuclei with >5% mitochondrial reads and those with unique feature counts over 10,000 or less than 200 using Seurat version 5 ^21^. All sn/scRNA-seq data were normalized via negative binomial regression corrected for mitochondrial counts using SCTransform ^21^. Cell types were identified using differentially expressed genes.

## Statistical Analyses

### Data Processing

Bulk RNA-seq samples underwent alignment to the respective genomes (hg19, mmul_10, mm10) using STAR aligner version 2.5.3.a (^22^; --outSAMstrandField intronMotif – outFilterScoreMinOverLread 0.3 --outFilterScoreMinOverRread 0.3 and –twopassMode Basic) after preprocessing with Trimmomatic version 0.39 (^23^; reads < 36 bp long, leading or trailing reads < Phred score of 3 and allowing a maximum of 2 mismatches per read). RNA-seq read alignment yielded an average of 76,181,104 reads in humans (s.d. = 29,529,191; M%Alignment = 86.26%), 34,551,920 reads in primates (s.d. = 8,202,258; M%Alignment = 79.71%), and 32,317,723 (s.d. = 6,271,112) reads in mice (M%Alignment = 88.90%). Aligned reads were counted using featureCounts ^24^.

### Differential Expression

DESeq2 was employed for differential expression analysis in humans, primates, and HDID mice. Differential expression analyses in humans controlled for a few standard covariates: sex, age, post-mortem index (PMI) and brain pH level. Primate expression analyses controlled for cohort, which accounted for age and sex; and mouse analyses adjusted for sex since they were matched on all other covariates. We estimated transcriptome-wide correlations across differential expression results – using only homologous genes across species (via biomaRt; ^25^).

### Single-cell deconvolution

The abundance of cell types from sn/scRNA-seq data were predicted by group and brain region from bulk RNA-seq datasets via CIBERSORT (100 permutations; ^26^). That is, predictions of cell types in bulk RNA-seq data were inferred by referencing sn/scRNA-seq datasets to predict whether brain cell type abundances were different in alcohol users versus controls.

### Gene Co-expression Network Analyses using WGCNA

We performed gene network modeling via weighted gene co-expression analyses (WGCNA; ^27^). First, we created signed hybrid WGCNA networks from human post-mortem brain tissue - separately for each brain region examined. Before WGCNA gene network construction, we filtered genes that had less than 10 read counts in more than 90% of the samples and performed variance-stabilizing transformations as per WGCNA creators’ recommendations (https://horvath.genetics.ucla.edu/html/CoexpressionNetwork/Rpackages/WGCNA/faq.html). WGCNA models included 18,341, 16,474, and 16,755 genes for the PFC, NAc, and CeA, respectively. We weighted correlations between genes by using soft thresholds (6, 10, and 18) that were elected to maximize fit and satisfy WGCNA modeling assumptions (all scale free topology > 0.851). With this weighted correlation matrix, we created an adjacency matrix and then a distance matrix, which was hierarchically clustered and split into discrete gene networks using a dynamic tree cutting algorithm. Gene networks were arbitrarily assigned a color (e.g. the magenta gene network in the PFC is different than the magenta network in the CeA).

### Conservation, Association, and Characterization of Gene Co-expression Networks

We assessed the reproducibility of human gene co-expression networks and their conservation in primate and mouse models using the modulePreservation function. This function determines the pair-wise concordance of gene-gene correlations within a gene co-expression network between input and test data ^28^. As recommended by Langfelder et al., 2011, we evaluated the Zsummary metric, which is a composite of 7 different preservation statistics. A Zsummary value > 10 represents strong reproducibility and was used as our threshold for conservation. First, we performed module preservation analyses on 100 bootstrapped human brain samples. All WGCNA networks had a Zsummary score > 10.65 indicating robust and stable gene co-expression networks.

Using an effect-size approach, we investigated whether WGCNA networks were associated with alcohol outcomes by permutating the average absolute value of differential expression test statistic of genes in gene co-expression network (10,000 permutations). Gene co-expression networks were tested for enrichment in KEGG pathways, biological processes, molecular functions, and cellular component ontological processes using enrichR ^29^ as well as human brain cell-types via cell-specific enrichment analysis (CSEA; ^30^). We characterized hub genes using two approaches: expression-based Phenome-Wide Association Studies (ePheWAS; https://www.systems-genetics.org/ephewas) and querying the NHGRI-EBI catalogue of human genome-wide association studies (GWAS; https://www.ebi.ac.uk/gwas/).

### Partitioned Heritability

We performed partitioned heritability analyses using Linkage Disequilibrium Score Regression (LDSC; ^31, 32^). Partitioned LDSC estimates whether single nucleotide polymorphisms (SNPs) in and around a gene-set account for a significant proportion of the heritability of a trait proportional to the amount of variants included. To determine behavioral specificity, we tested whether the individual heritability of multiple human alcohol and tobacco use traits were enriched for all genes within conserved WGCNA networks associated with human and animal chronic alcohol use (across all brain regions). Our analyses used LDSC’s default gene window size (100 kb) to assign SNPs to genes based on the expectation that up to 80% of local gene regulatory regions occur within 100 kb of a gene ^33^.

## RESULTS

Using transcriptome-wide samples of 92 humans, 53 primates, and 90 mice in three brain regions (see Table 1), we assessed the overlap of AUD RNA signature with pre-clinical chronic alcohol use. Using PFC, NAc, and CeA RNA-seq data on a sample of 60 humans, our analyses revealed significant correlations between differential expression results of human AUD and mammalian models of alcohol use (see Figure 1A; all p_adj_ < 0.05). Cross-species correlations differed by tissue, species, and preclinical model. The most significant and positive correlations with human AUD occurred in primates and mouse models of binge drinking. Some correlations were negative. For instance, mouse chronic intermittent ethanol exposure (CIE) and alcohol self-administration data exhibited significant negative correlations with AUD in the PFC, but these models showed no significant correlations with AUD in the NAc or CeA. It is important to note that regional specificity in brain regions across species are not completely identical.

**Figure 1.**
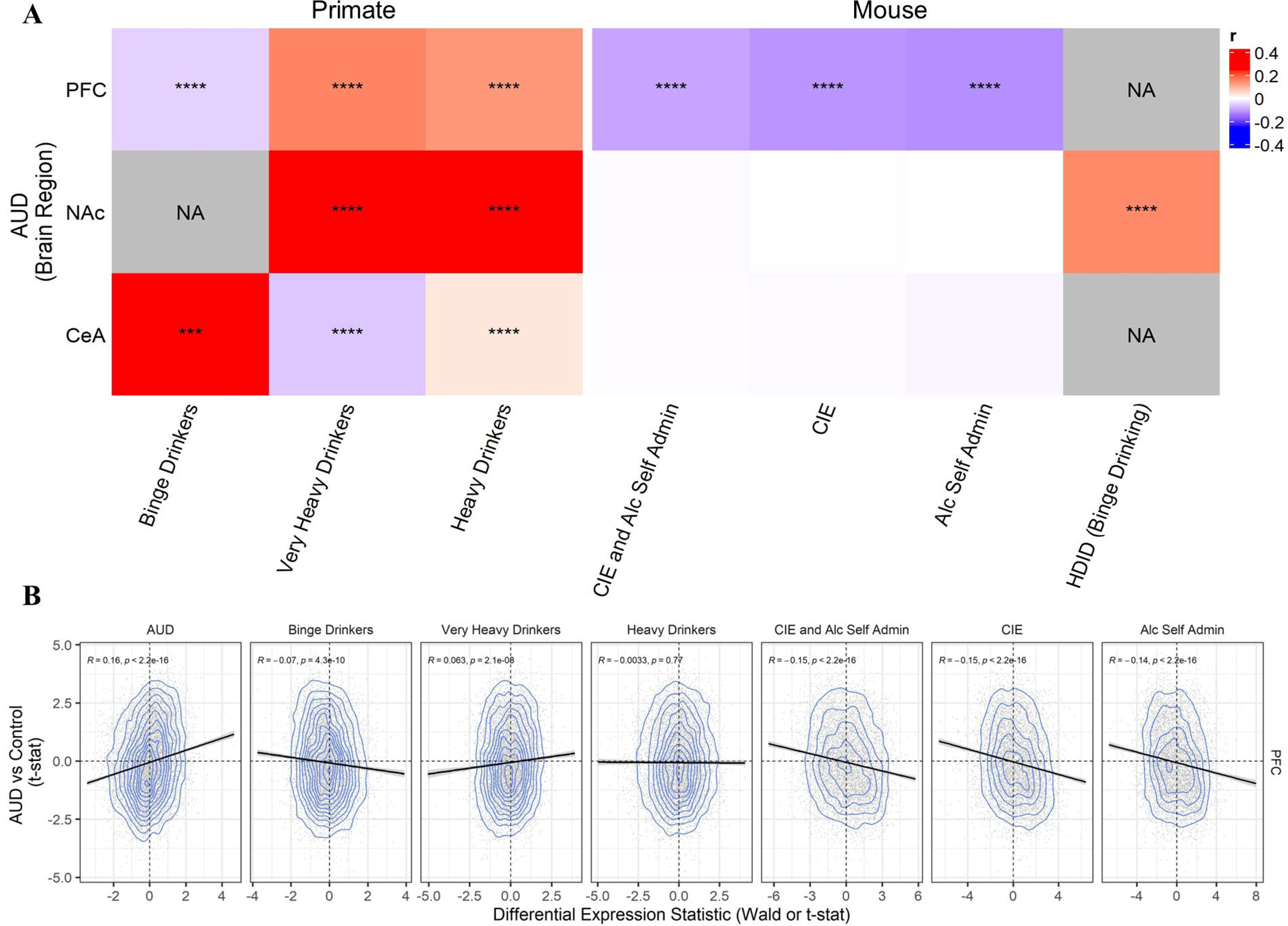
AUD Brain RNA Signature Robustly Correlated with Animal Models of Alcohol Use. Transcriptome-wide correlations of differentially expressed genes across traits, brain regions and species. A) Heatmap of AUD RNA correlations with mouse and primate models of alcohol use; **** p < 0.001; blank = p > 0.05; NA = not applicable B) Scatterplot showing reproducibility of brain RNA correlations using an independent sample of AUD; R = Pearson correlation coefficient. Self Admin = ethanol self-administration; CIE = chronic intermittent ethanol exposure; HDID = high drinking in the dark; PFC = pre-frontal cortex, NAc = nucleus accumbens; CeA = central nucleus of amygdala.

To examine the reproducibility of RNA correlations, we used an independent microarray sample of 32 individuals with AUD and matched controls in the PFC ^11^. AUD RNA signatures correlated in the expected direction across samples, although with a small effect size (r = 0.15, p < 0.0001). All but the heavy drinking primate model showed significant correlations with AUD (see Figure 1B), suggesting our findings are reproducible. Notably, the very heavy drinking primate model showed correlations that aligned with human AUD at a level comparable to 39% of the correlation observed relative to the human AUD correlation with itself across different samples (ie., r = 0.16 for AUD-AUD; r = 0.063 for VHD-AUD; see Figure 1B).

Next, we used publicly available snRNA-sequencing data from post-mortem human and fetal primate PFC (11,196-12,803 cells after QC), striatum (12,243-19,862 cells after QC) and amygdala (13,422-13,984 cells after QC) as well as mouse NAc (41,519 cells after QC) studies to investigate the cellular basis of AUD and mammalian alcohol use. Only bulk RNA-seq data and not microarray data (i.e., CIE and alcohol self-administration) were integrated with snRNA-seq data to avoid methodological biases. Note that many, but not all, brain cell-types were similar across species (see Figure 2). Using single-cell deconvolution techniques (CIBERSORT), we identified that the proportion of oligodendrocytes was decreased in human AUD vs controls in the PFC and the NAc (see Figure 2). Similarly, the proportion of oligodendrocytes were decreased in the NAc of a mouse model of binge drinking (high drinking in the dark [HDID]; ^15^) and oligodendrocyte progenitor cells were decreased in the PFC of primate models of alcohol use (all P < 0.05; see Figure 2). A few other trends were observed in AUD and showed similar effects in primates and mice. For instance, the relative proportion of NAc dopamine receptor 2 medium spiny neurons (MSN_D2) were trending to decrease in AUD and were decreased in HDID binge drinking mice. Additionally, the relative proportion of CeA excitatory neurons were trending to be decreased in AUD and were also decreased in binge drinking primates. Reassuringly, the directionality was consistent across species and some proportion of cell-types showed a dose-response, but the lack of significance could reflect low statistical power.

**Figure 2.**
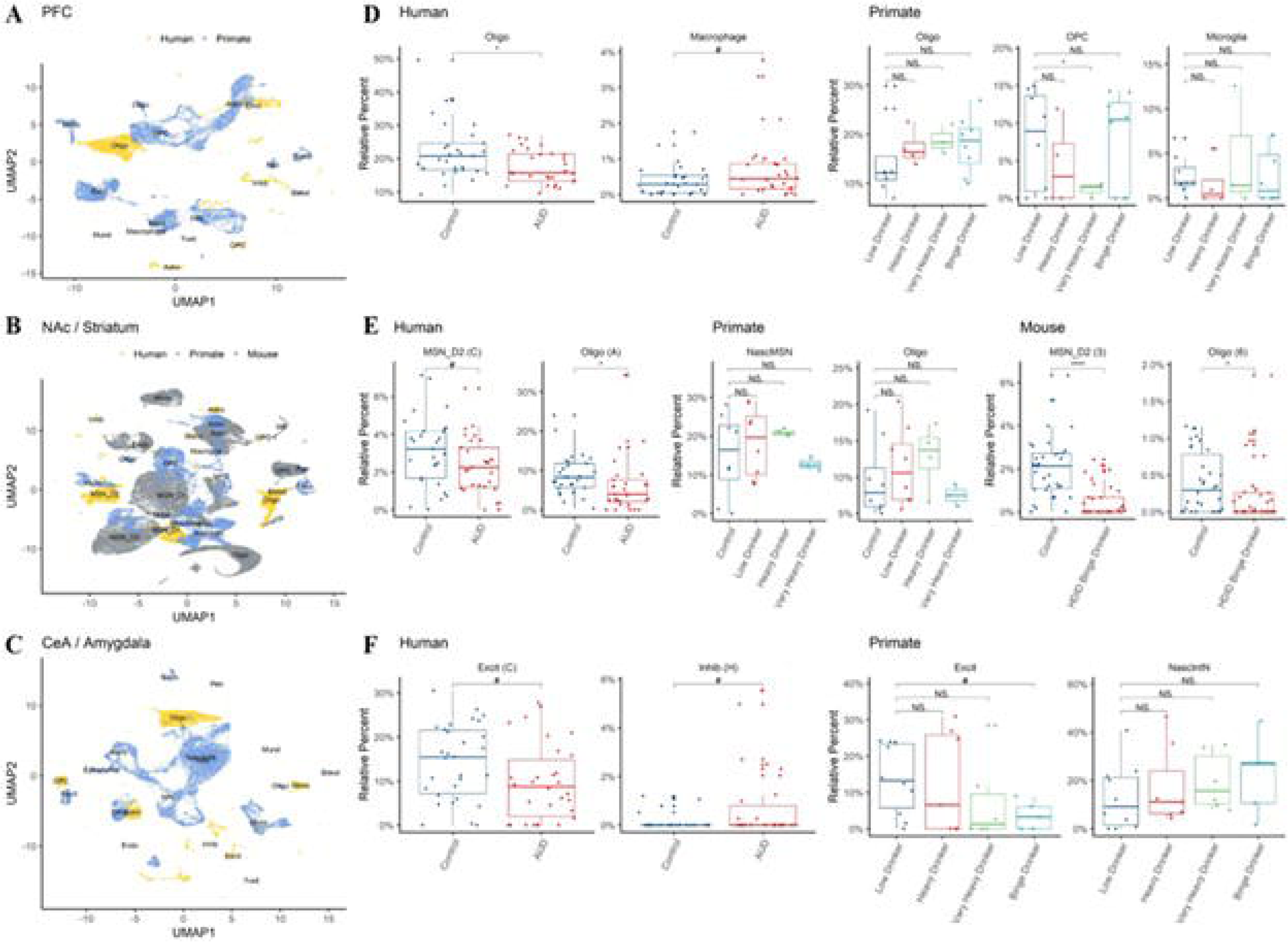
Single-cell Deconvolution of AUD and Alcohol Use Brain Signatures Across Animal Models and Brain Regions. In silico cytometry of human AUD and preclinical alcohol use across traits, brain regions, and species. A-C shows clustering of cell-types by brain region and species. D-F shows the proportion of each cell type by group across species, brain regions and pre-clinical models. HDID = high drinking in the dark; Oligo = oligodendrocytes, Excit = excitatory neuron; Inhib = inhibitory neuron; NascIntN = nascent interneuron; MSN_D2 = dopamine D2 receptor medium spiny neuron; OPC = oligodendrocyte progenitor cell; PFC = pre-frontal cortex, NAc = nucleus accumbens; CeA = central nucleus of amygdala; **** p < 0.001; * p < 0.05; # p < 0.1

Next, using the 60 RNA-seq samples of AUD and matched controls, we created /gene co-expression networks via weighted gene co-expression network analyses (WGCNA) gene in each brain region individually and tested for their conservation across species. WGCNA uses pairwise RNA correlations of all gene pairs, clusters similar correlated gene pairs together and creates distinct gene co-expression networks arbitrarily assigned to a color. We tested whether human gene co-expression networks were conserved across species (and within brain region) using permutation analyses and a recommended threshold outlined in prior work (Z_summary_ > 10; ^28^; see Supplementary Methods). We found that 6, 16, and 8 networks were highly conserved across species in the PFC, NAc and CeA, respectively (Z_summary_ > 10; see Supplementary Figure S1). Of the 30 highly conserved WGCNA networks across species, 80% were highly conserved in primates and 46.67% were highly conserved in mice. Eight conserved gene co-expression networks were associated with both human AUD samples and at least one alcohol use outcome in animals (see Figure 3). All gene co-expression networks were associated with AUD and mammalian alcohol traits in the same direction. All gene co-expression networks were robustly enriched for certain brain cell types and all but one was significantly enriched for gene ontology functions (see Table 2).

**Figure 3.**
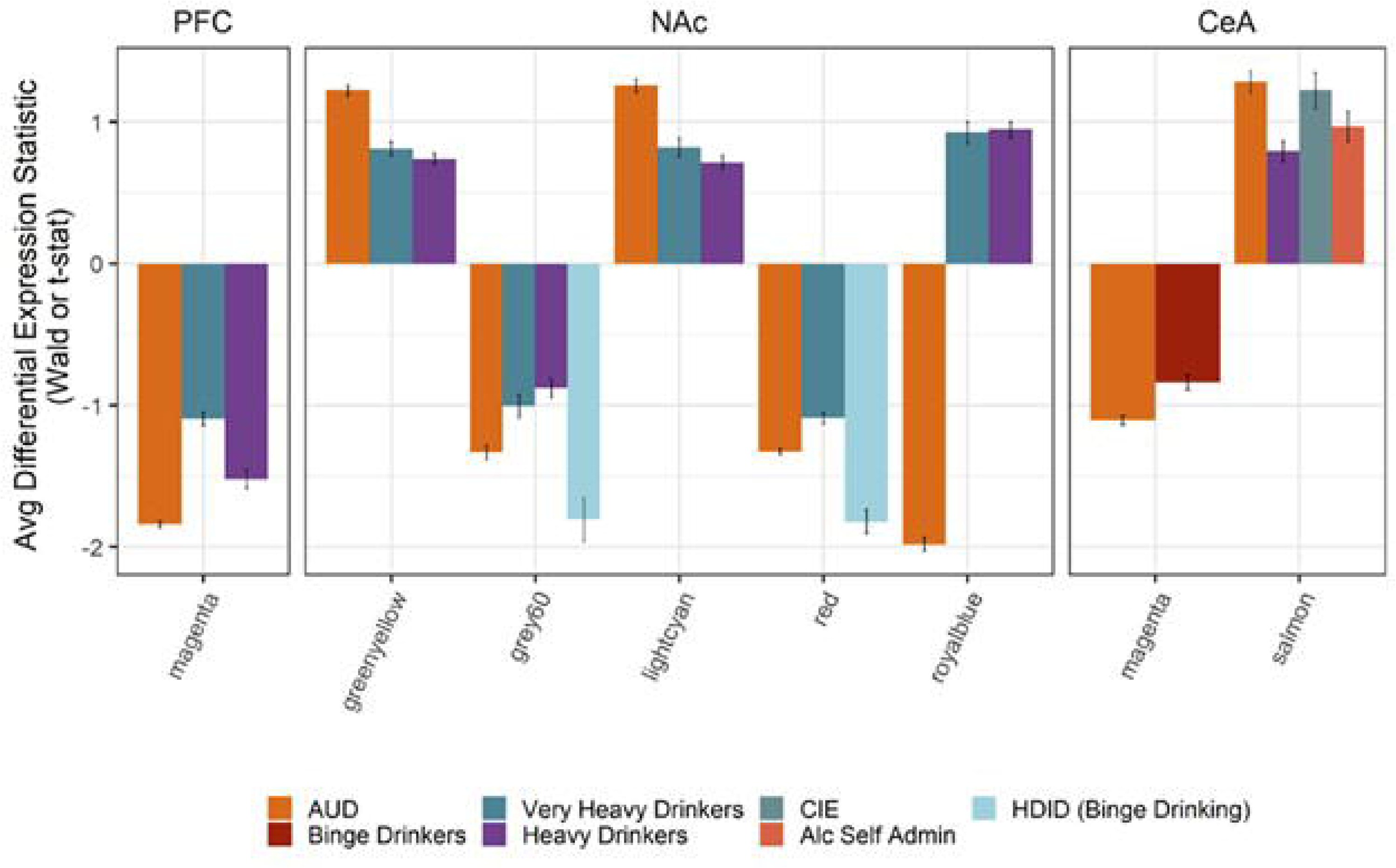
Conserved Gene Co-expression Networks Associated with Human AUD and Animal Models of Alcohol Use. Gene co-expression networks associated with alcohol use across samples and species. X-axis shows the WGCNA gene co-expression network and the y-axis shows the mean differential expression test statistic by brain region and trait. All gene networks in this plot were significant via permutation analyses (padj < 0.05). WGCNA = weighted gene co-expression network analysis; CIE = chronic intermittent alcohol exposure; HDID = high drinking in the dark. Note color for WGCNA networks are arbitrary and the magenta gene network in the PFC is distinct from the magenta gene network in the CeA.

**Table 2.** WGCNA Gene Networks were Conserved Across Species and Associated with Human AUD and Chronic Alcohol Use.

| *Cell Specific Enrichment Analyses of WGCNA Networks* |  |  |  |  |
| --- | --- | --- | --- | --- |
| Gene Network | Brain Region | Most Enriched Cell Type | P | Padj |
| Magenta | PFC | Myelinating Oligodendrocytes | 2.12E-109 | 3.72E-108 |
| Greenyellow | NAc | Bergman Glia & Mature Oligodendrocytes | 9.59E-11 | 3.36E-09 |
| Grey60 | NAc | Drd2+ Medium Spiny Neurons | 5.155e-12 | 1.80E-10 |
| Lightcyan | NAc | Oligodendrocyte Progenitor Cells | 2.378e-06 | 2.08E-05 |
| Red | NAc | Drd1+ Medium Spiny Neurons | 1.54E-39 | 2.69E-38 |
| Royalblue | NAc | Cholinergic Neurons | 1.243e-08 | 4.35E-07 |
| Magenta | Ce A | Myelinating Oligodendrocytes | 2.90E-81 | 1.02E-79 |
| Salmon | Ce A | Immune Cells | 2.837e-11 | 9.93E-10 |

*Gene Ontology Enrichment Analyses of WGCNA Networks*
| Gene Network | Brain Region | Most Enriched Ontological Process | P | Padj |
| --- | --- | --- | --- | --- |
| Magenta | PFC | Myelination (BP) | 2.242E-06 | 0.004556 |
| Greenyellow | NAc | Macrophage cytokine production (BP) | 4.97E-07 | 0.001042 |
| Grey60 | NAc | Regulation of neurological systems (BP) | 0.001468 | 0.2105 |
| Lightcyan | NAc | Central nervous system development (BP) | 5.877E-05 | 0.02891 |
| Red | NAc | Anterograde trans-synaptic signaling (BP) | 5.68E-14 | 1.27E-10 |
| Royalblue | NAc | Gap junction (KEGG) | 5.134E-05 | 0.008677 |
| Magenta | CeA | Actin cytoskeleton (CC) | 0.000194 | 0.01855 |
| Salmon | CeA | Staphylococcus aureus infection (KEGG Pathway) | 6.39E-25 | 1.19E-22 |

Note only the most significant enrichment is shown. Only the Grey60 gene network was not significantly enriched for any gene ontology process. WGCNA= weighted gene co-expression analyses; sPFC = Superior Pre-Frontal Cortex; NAc = Nucleus Accumbens; Ce A = Central Nucleus of the Amygdala; fBP = biological process; KEGG = Kyoto encyclopedia of genes and genomes; CC = cellular component. Note color for WGCNA networks are arbitrary and the magenta gene network in the PFC is distinct from the magenta gene network in the CeA

To identify key regulators that may be driving these physiological processes, we sought to find and characterize hub genes in these WGCNA gene co-expression networks. Selecting the top 2.5% of module membership in each gene network, we found a total of 63 hub genes (see Figure 4; 57 unique genes). We characterized these genes with two online tools: NHGRI and ePheWAS and reported these results in Table 3. In short, we found these genes may be implicated in various drug use traits, as well as impulsivity, motivation, stress, and other molecular and morphological outcomes.

**Figure 4.**
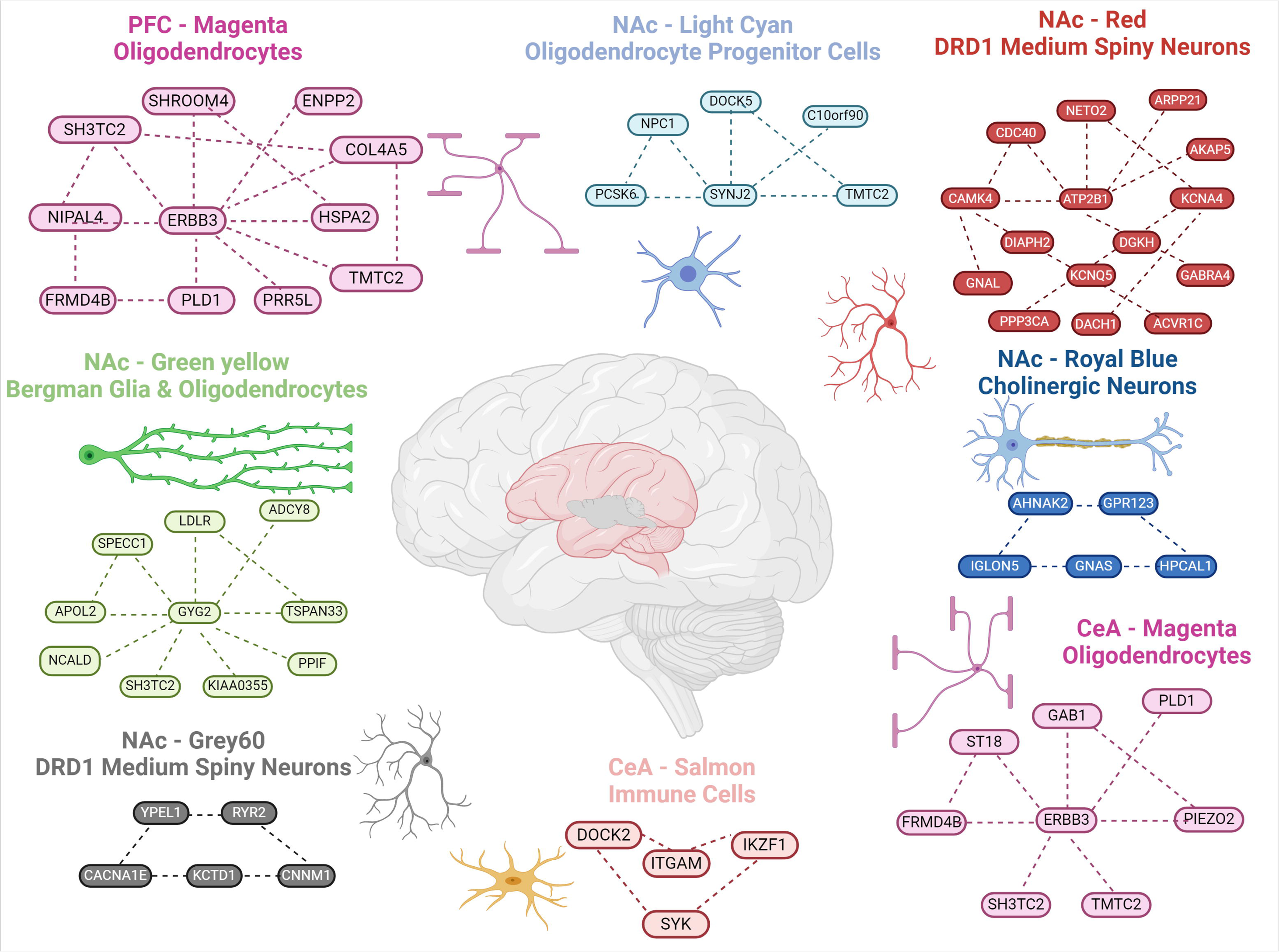
Conserved Gene Co-expression Networks Associated with Human AUD and Animal Models of Alcohol use. Image created via Biorender.com. Note enriched cell-types are displayed next to WGCNA networks and dashed lines between genes are for visual purposes only and not derived from gene-gene connectivity metrics. Bottom right two gene networks are in the CeA, the top left is in the PFC and the rest are in the NAc.

**Table 3.**
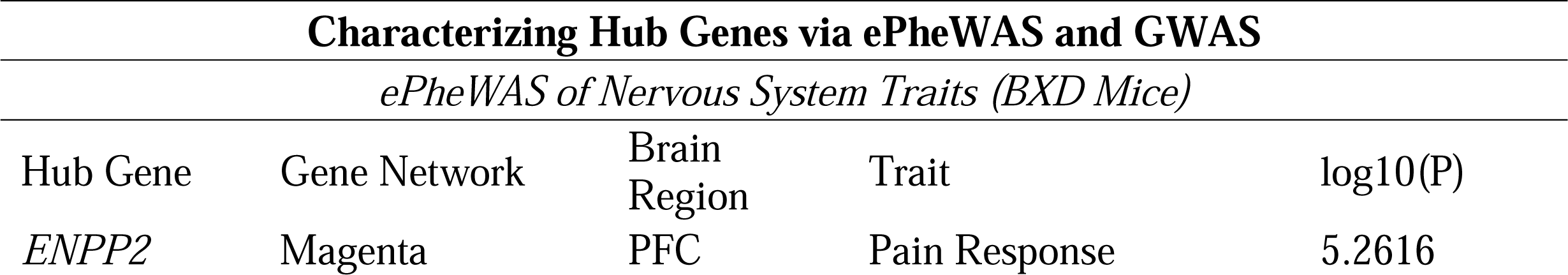
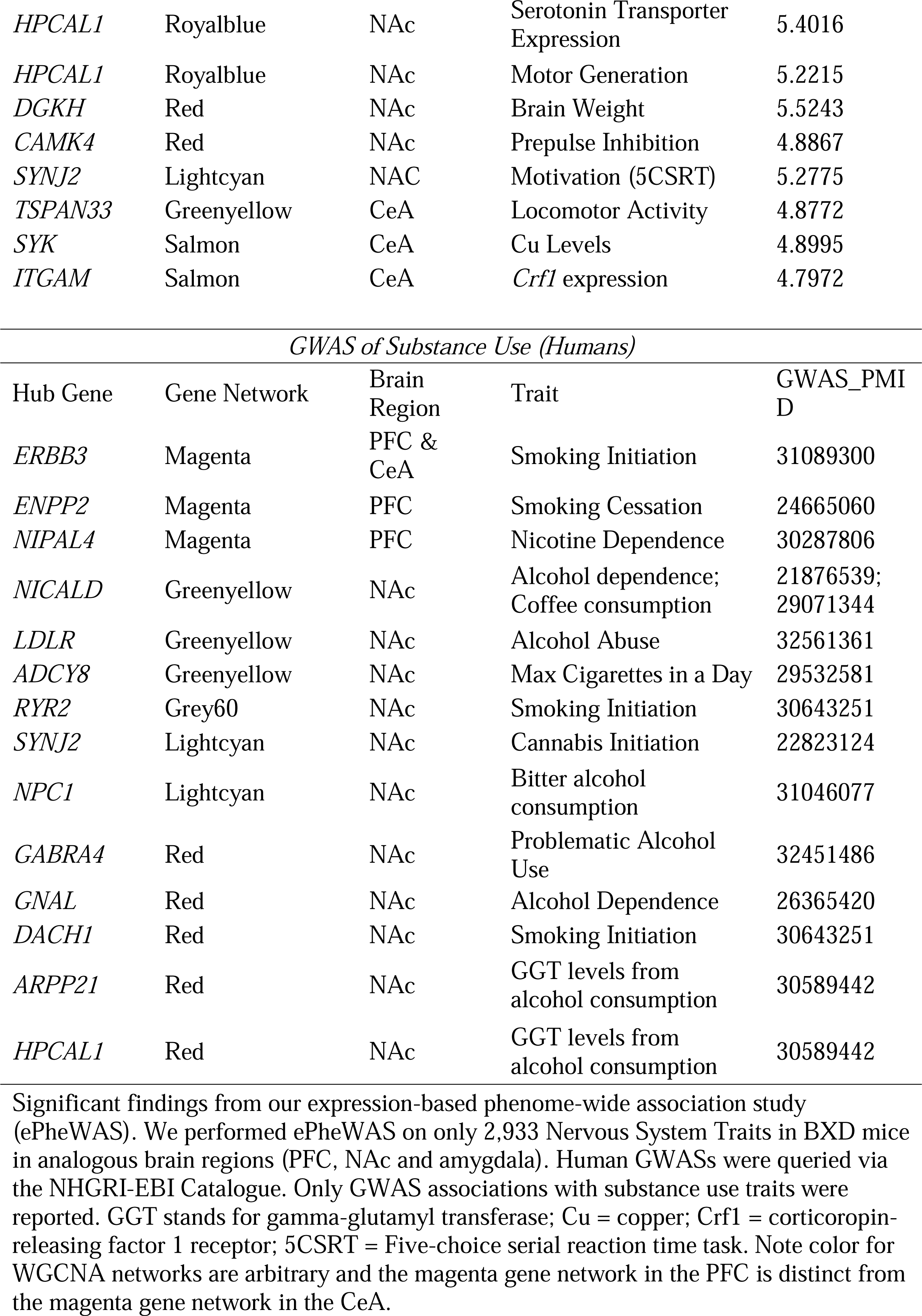
Hub genes from WGCNA Gene Networks that were Conserved Across Species and Associated with Human AUD and Animal Chronic Alcohol Use.

Lastly, we used partitioned heritability analyses to test for genetic enrichment. The heritability of drinks per week and problematic alcohol use, but not tobacco nor a non-substance use trait (wears glasses), were significantly enriched for the genes from the conserved gene co-expression networks associated with AUD and alcohol use (see Figure 5).

**Figure 5.**
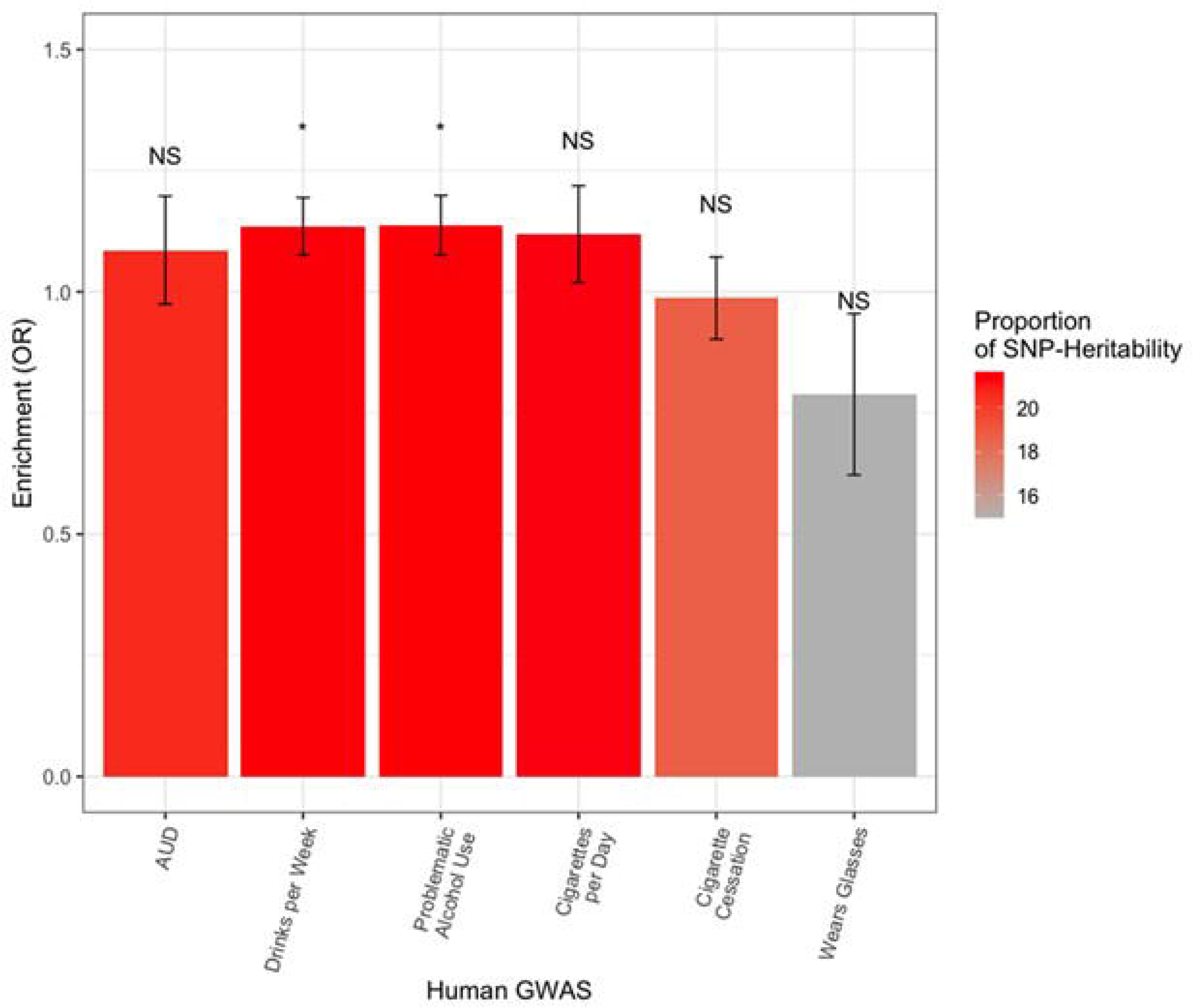
Conserved AUD & Alcohol Use Networks Enriched for GWAS Associations of Alcohol Use. Results from partitioned heritability analyses. DNA variants within 100kb of the genes from conserved WGCNA networks that were associated with human AUD and mammalian models of alcohol use were included in these analyses. * P < 0.05; NS = P > 0.05.

## DISCUSSION

This study found partial support for our hypotheses. RNA expression and cell-type abundances associated with human AUD generally correlated with animal models of alcohol use, but the extent of which depended on species, brain region, and behavioral trait. The strongest overlap of brain RNA patterns was observed between humans and primates, which matches the close phylogenetic relationship between species. Mouse binge drinking models showed substantial overlap with human AUD gene expression, which is consistent with independent biological data on human and mouse alcohol use phenotypes ^34^. Importantly, brain correlations with primate and mouse alcohol use were reproducible with an independent human AUD sample ^11^ highlighting the reproducibility of these findings.

Single-cell transcriptomic data provided insights into the cellular basis of AUD and mammalian chronic alcohol use. The abundance of oligodendrocyte cell types were decreased in AUD among humans and in the animal models of alcohol use in the PFC and NAc. These observations are consistent with snRNA-seq data in AUD ^35^, as well as postmortem cellular and in vivo neuroimaging data, suggesting that prolonged alcohol use is linked to oligodendrocyte and white-matter damage ^36^. We also observed parallel trends suggesting decreased abundance of dopamine MSN_D2 NAc medium spiny neurons (MSN) in human AUD and mouse binge drinking. D2 MSNs in NAc are thought to mediate reward, motivation, and addictive behaviors. Inhibiting NAc MSN D2 neurons has been shown to experimentally promote alcohol intake ^35^. While these analyses had limited statistical power, it was reassuring to detect consistent directionality across species and mirror similar observations in the literature.

Utilizing gene co-expression network analyses, we were able to identify conserved systems across species. These analyses suggested that dopamine and acetylcholine neurons, as well as a variety of glial cells, play a role in cross-species consilience in various brain regions and alcohol use traits. Pathways and gene-sets involved in the inflammation, myelination and synaptic plasticity were enriched among these gene-networks. Expression of the hub genes from these gene networks were linked with relevant traits in mice via ePheWAS: motivation, impulsivity and locomotor response [1] [2], as well as pertinent traits via human GWASs: alcohol, tobacco and cannabis use. Accordingly, genes from these co-expression networks accounted for ∼20% of the heritability of alcohol use traits but were not enriched in a negative control trait (wearing glasses). In sum, despite the challenges inherent to harmonizing such disparate data sources, there is significant face validity and coherence with known addiction biology to suggest that the approach used here has utility.

## CONCLUSION

To our knowledge, this is the first biologically relevant study on alcohol use to aggregate across monkeys, mice, and (hu)mankind. The heterogeneity across studies complicates interpretations. For instance, we are unable to decipher the degree to which an association – or lack thereof – is due to the degree of alcohol exposure, genetics, mis-match of brain regions or technical factors. Sometimes associations of differentially expressed genes across species were negative, which is in line with previous cross-species research on drug use traits but is not understood well (^34^; ^37^). Data used in the current paper lacks diversity in genetics and sex and may limit the generalizability of our findings. Cell-type abundance analyses should be interpreted with caution in light of small (underpowered) samples, containing few cells and sometimes using less relevant samples (i.e., fetal primate data). Notwithstanding the limitations, our paper provides a data-driven approach to highlight the biological intersection between human AUD and animal alcohol use and systematically compares pre-clinical paradigms of alcohol use and their relevance to human AUD. Future efforts would benefit from harmonizing data with larger samples, and more traits/substances and incorporating different techniques to synergistically assess the biological overlap across species, cell types, and brain regions/subregions.

## Disclaimer

The findings and conclusions in this publication are those of the authors, and do not represent the views of the U.S. Department of Veterans Affairs, the U.S. Department of Agriculture, and do not represent any US Government determination, position, or policy.

## Acknowledgements

We would like to thank prior human and animal researchers for making this study possible through their engagement in data sharing on public platforms.

## References

1. Kranzler HR, Zhou H, Kember RL, Vickers Smith R, Justice AC, Damrauer S et al. Genome-wide association study of alcohol consumption and use disorder in 274,424 individuals from multiple populations. Nat Commun 2019; 10(1): 1499.

2. Koob GF, Volkow ND. Neurobiology of addiction: a neurocircuitry analysis. The lancet Psychiatry 2016; 3(8): 760–773.

3. Palmer RHC, Johnson EC, Won H, Polimanti R, Kapoor M, Chitre A et al. Integration of evidence across human and model organism studies: A meeting report. Genes Brain Behav 2021; 20(6): e12738.

4. Becker HC, Lopez MF. An Animal Model of Alcohol Dependence to Screen Medications for Treating Alcoholism. Int Rev Neurobiol 2016; 126: 157–177.

5. Vena AA, Zandy SL, Cofresi RU, Gonzales RA. Behavioral, neurobiological, and neurochemical mechanisms of ethanol self-administration: A translational review. Pharmacol Ther 2020; 212: 107573.

6. Monleon S, Duque A, Mesa-Gresa P, Redolat R, Vinader-Caerols C. An Animal Model of Alcohol Binge Drinking: Chronic-Intermittent Ethanol Administration in Rodents. Methods Mol Biol 2019; 2011: 281–293.

7. Preuss TM, Wise SP. Evolution of prefrontal cortex. Neuropsychopharmacology 2022; 47(1): 3–19.

8. Barton RA, Harvey PH. Mosaic evolution of brain structure in mammals. Nature 2000; 405(6790): 1055-1058.

9. Lebish IJ, Moraski RM. Mechanisms of immunomodulation by drugs. Toxicol Pathol 1987; 15(3): 338–345.

10. Rao X, Thapa KS, Chen AB, Lin H, Gao H, Reiter JL et al. Allele-specific expression and high-throughput reporter assay reveal functional genetic variants associated with alcohol use disorders. Mol Psychiatry 2021; 26(4): 1142–1151.

11. Ponomarev I, Wang S, Zhang L, Harris RA, Mayfield RD. Gene coexpression networks in human brain identify epigenetic modifications in alcohol dependence. J Neurosci 2012; 32(5): 1884–1897.

12. Iancu OD, Colville A, Walter NAR, Darakjian P, Oberbeck DL, Daunais JB et al. On the relationships in rhesus macaques between chronic ethanol consumption and the brain transcriptome. Addict Biol 2018; 23(1): 196–205.

13. Walter N, Cervera-Juanes R, Zheng C, Darakjian P, Lockwood D, Cuzon-Carlson V et al. Effect of chronic ethanol consumption in rhesus macaques on the nucleus accumbens core transcriptome. Addict Biol 2021; 26(5): e13021.

14. Baker EJ, Farro J, Gonzales S, Helms C, Grant KA. Chronic alcohol self-administration in monkeys shows long-term quantity/frequency categorical stability. Alcohol Clin Exp Res 2014; 38(11): 2835–2843.

15. Hitzemann R, Oberbeck D, Iancu O, Darakjian P, McWeeney S, Spence S et al. Alignment of the transcriptome with individual variation in animals selectively bred for High Drinking-In-the-Dark (HDID). Alcohol 2017; 60: 115–120.

16. Smith ML, Lopez MF, Wolen AR, Becker HC, Miles MF. Brain regional gene expression network analysis identifies unique interactions between chronic ethanol exposure and consumption. PLoS One 2020; 15(5): e0233319.

17. Goldstein DB. Relationship of alcohol dose to intensity of withdrawal signs in mice. J Pharmacol Exp Ther 1972; 180(2): 203–215.

18. Tran MN, Maynard KR, Spangler A, Huuki LA, Montgomery KD, Sadashivaiah V et al. Single-nucleus transcriptome analysis reveals cell-type-specific molecular signatures across reward circuitry in the human brain. Neuron 2021; 109(19): 3088–3103 e3085.

19. Zhu Y, Sousa AMM, Gao T, Skarica M, Li M, Santpere G et al. Spatiotemporal transcriptomic divergence across human and macaque brain development. Science 2018; 362(6420).

20. Chen R, Blosser TR, Djekidel MN, Hao J, Bhattacherjee A, Chen W et al. Decoding molecular and cellular heterogeneity of mouse nucleus accumbens. Nat Neurosci 2021; 24(12): 1757–1771.

21. Hao Y, Stuart T, Kowalski MH, Choudhary S, Hoffman P, Hartman A et al. Dictionary learning for integrative, multimodal and scalable single-cell analysis. Nat Biotechnol 2024; 42(2): 293–304.

22. Dobin A, Davis CA, Schlesinger F, Drenkow J, Zaleski C, Jha S et al. STAR: ultrafast universal RNA-seq aligner. Bioinformatics 2013; 29(1): 15–21.

23. Bolger AM, Lohse M, Usadel B. Trimmomatic: a flexible trimmer for Illumina sequence data. Bioinformatics 2014; 30(15): 2114–2120.

24. Liao Y, Smyth GK, Shi W. featureCounts: an efficient general purpose program for assigning sequence reads to genomic features. Bioinformatics 2014; 30(7): 923–930.

25. Smedley D, Haider S, Ballester B, Holland R, London D, Thorisson G et al. BioMart--biological queries made easy. BMC Genomics 2009; 10: 22.

26. Newman AM, Liu CL, Green MR, Gentles AJ, Feng W, Xu Y et al. Robust enumeration of cell subsets from tissue expression profiles. Nat Methods 2015; 12(5): 453–457.

27. Langfelder P, Horvath S. WGCNA: an R package for weighted correlation network analysis. BMC Bioinformatics 2008; 9: 559.

28. Langfelder P, Luo R, Oldham MC, Horvath S. Is my network module preserved and reproducible? PLoS Comput Biol 2011; 7(1): e1001057.

29. Kuleshov MV, Jones MR, Rouillard AD, Fernandez NF, Duan Q, Wang Z et al. Enrichr: a comprehensive gene set enrichment analysis web server 2016 update. Nucleic acids research 2016; 44(W1): W90–97.

30. Lambert J, Wijermans PW, Dekker GA, Ossenkoppele GJ. Chemotherapy in non-Hodgkin’s lymphoma during pregnancy. Neth J Med 1991; 38(1-2): 80–85.

31. Finucane HK, Bulik-Sullivan B, Gusev A, Trynka G, Reshef Y, Loh PR et al. Partitioning heritability by functional annotation using genome-wide association summary statistics. Nat Genet 2015; 47(11): 1228–1235.

32. Bulik-Sullivan BK, Loh PR, Finucane HK, Ripke S, Yang J, Schizophrenia Working Group of the Psychiatric Genomics C et al. LD Score regression distinguishes confounding from polygenicity in genome-wide association studies. Nat Genet 2015; 47(3): 291–295.

33. Gandal MJ, Zhang P, Hadjimichael E, Walker RL, Chen C, Liu S et al. Transcriptome-wide isoform-level dysregulation in ASD, schizophrenia, and bipolar disorder. Science 2018; 362(6420).

34. Huggett SB, Johnson EC, Hatoum AS, Lai D, Srijeyanthan J, Bubier JA et al. Genes identified in rodent studies of alcohol intake are enriched for heritability of human substance use. Alcoholism, clinical and experimental research 2021; 45(12): 2485–2494.

35. van den Oord E, Xie LY, Zhao M, Aberg KA, Clark SL. A single-nucleus transcriptomics study of alcohol use disorder in the nucleus accumbens. Addict Biol 2023; 28(1): e13250.

36. Miguel-Hidalgo JJ. Molecular Neuropathology of Astrocytes and Oligodendrocytes in Alcohol Use Disorders. Front Mol Neurosci 2018; 11: 78.

37. Enoch MA, Zhou Z, Kimura M, Mash DC, Yuan Q, Goldman D. GABAergic gene expression in postmortem hippocampus from alcoholics and cocaine addicts; corresponding findings in alcohol-naive P and NP rats. PLoS One 2012; 7(1): e29369.

