## Supplement for "Bulk and Single-cell Transcriptomic Brain Data Identify Overlapping Processes and Cell-types with Human AUD and Mammalian Models of Alcohol Use"

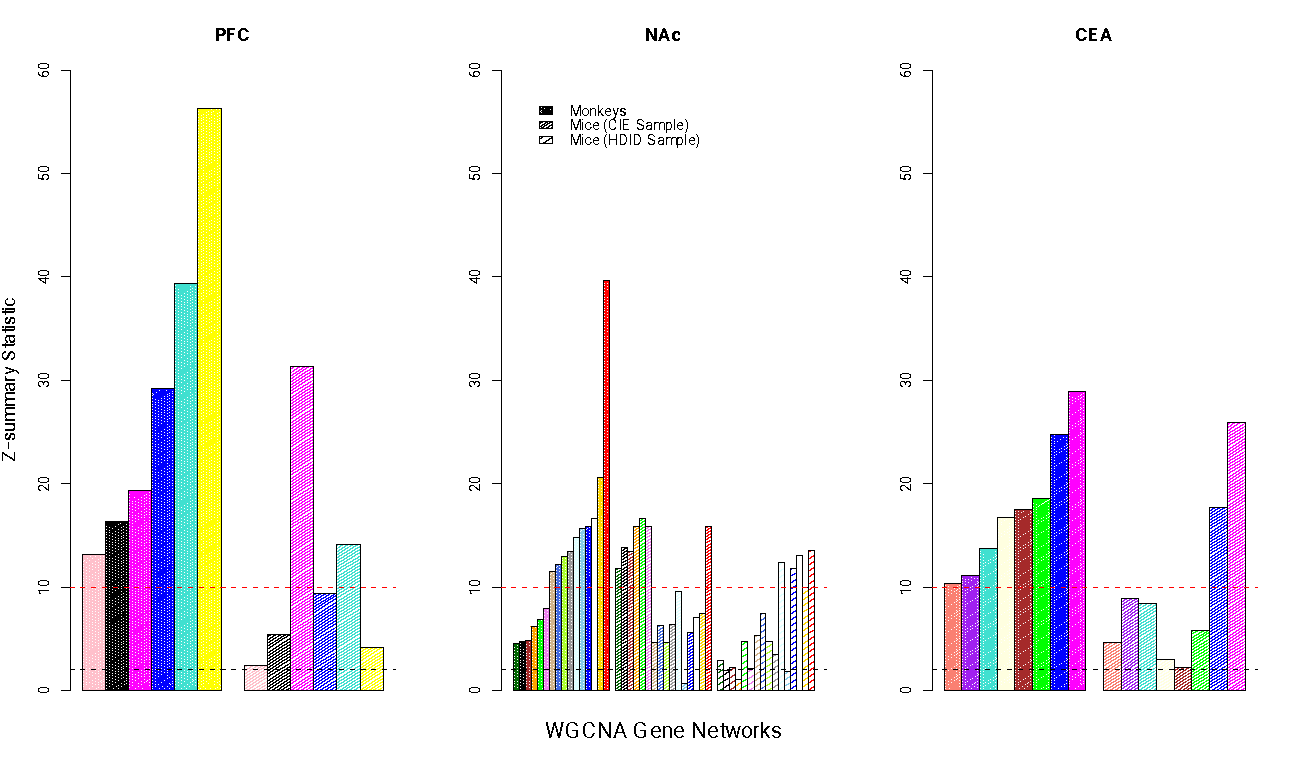

Gene network conservation across human AUD samples with primate and mouse models of alcohol use. Gene networks were made in human brain tissue and tested for their reproducibility in mammalian models of alcohol use. Zsummary is the metric of conservation. Zsummary > 10 indicates a highly reproducible gene co-expression network, a value between 2-10 would be weakly to moderately conserved and below 2 would not be conserved. WGCNA = weighted gene co-expression network analysis; CIE = chronic intermittent alcohol exposure; HDID = high drinking in the dark. Note color for WGCNA networks are arbitrary and the magenta gene network in the PFC is distinct from the magenta gene network in the CeA.
